# Host preference and survivorship of *Euschistus heros* (Hemiptera: Pentatomidae) strains on cotton and soybean

**DOI:** 10.1101/2022.06.04.494803

**Authors:** Frederico Hickmann, Erick M. G. Cordeiro, Mateus Souza L. Aurélio, Alan Valdir Saldanha, Alberto Soares Corrêa

## Abstract

The Neotropical brown stink bug *Euschistus heros* (Fabricius) (Hemiptera: Pentatomidae) is a key pest of soybeans, *Glycine max*, and recently became an economically important pest of cotton, *Gossypium hirsutum*. This stink bug has two allopatric strains, one prevalent in southern Brazil (SS), and another in the north (NS). The two strains hybridize in central Brazil. Knowledge of host preferences and host suitability of these strains can clarify the contribution of the different gene pools to contemporary adaptive features such as the ability to harm cotton crops. We tested the attraction of the *E. heros* strains and reciprocal hybrids [♀N × ♂S (HNS) and ♀S × ♂N (HSN)] to soybean and cotton plants and evaluated the nymph development and survivorship of the two strains and reciprocal hybrids fed on soybean or cotton. We conducted host-choice experiments with 4th instars and adult females and evaluated the survival of immatures on soybean and cotton plants in laboratory conditions. The SS strain preferred soybean over cotton. NS and hybrid strains chose randomly between soybean and cotton plants. All strains developed on soybean, with similar survival rates. On cotton, the pure strains did not reach adulthood; however, the hybrids developed on cotton but with a survival rate less than 1%. Our results showed that *E. heros* SS was more attracted to soybeans, and NS and hybrid strains had a polyphagous choice behavior, suggesting that current host selection has been mediated by historical and, mainly, contemporary relationships of *E. heros* strains with these hosts.

## 1. Introduction

Herbivorous insects have a close relationship to their host plants and are subject to continuous variation over time (Simon et al., 2015). Expansion of agricultural frontiers dramatically changes the landscape by replacing complex natural areas with simplified agroecosystems (Silva et al., 2020). Insect populations exploiting crops encounter large patches of single hosts, resulting in diet specialization or evolution of novel plant-insect interactions (Jaenike, 1990; Wetzel et al., 2016). Therefore, agroecosystems are important areas for evolutionary change (especially in superabundant monocultures) and may lead to rapid selection for host use in populations of crop pests [e.g., corn leafhopper, *Dalbulus maidis* (DeLong & Wolcott); western corn rootworm, *Diabrotica virgifera virgifera* LeConte; and the cotton fleahopper, *Pseudatomoscelis seriatus* (Reuter)] (Barman et al., 2012; Bernal and Medina, 2018; Dávila-Flores et al., 2013; Gray et al., 2009; Medina et al., 2012).

The Neotropical brown stink bug, *Euschistus heros* (Fabricius) (Hemiptera: Pentatomidae), is a polyphagous pest recorded on 21 plant species, and can complete its life cycle on six, primarily Fabaceae (Smaniotto and Panizzi, 2015). Considered in the 1980s as a secondary pest on soybeans, *Glycine max* (L.) Merrill, currently *E. heros* is a serious pest that feeds on soybean pods, causing them to abort and reducing grain weight and quality (Panizzi et al., 2000; Panizzi and Slansky, 1985; Sosa-Gómez et al., 2020). Furthermore, in the last decade, reports of *E. heros* moving to cotton crops, *Gossypium hirsutum* L., after soybean harvest have become frequent, mainly in the central region of Brazil (Soria et al., 2017, 2011, 2010, 2009). This behavior has been hypothesized to be a host range expansion, since an increasing number of field reports indicate that *E. heros* increases annually in cotton crops in Brazil.

Two allopatric strains of *E. heros* occur in Brazil, one prevalent in the south (hereafter SS) and another common in the north/northeast (hereafter NS) (Soares et al., 2018). Despite the long period of geographic isolation between the northern and southern strains, these allopatric strains have formed a hybridization zone in central Brazil, without significant reproductive isolation (Hickmann et al., 2021; Soares et al., 2018; Zucchi et al., 2019). The reunion of two gene pools, without complete reproductive isolation, may mediate evolutionary events through recombination, hybrid vigor, and selection of adaptive alleles in local populations (Corrêa et al., 2019; Gasperi et al., 1991; Wilding et al., 2001).

In Brazil, cotton is grown mainly in the central part of the country, which coincides with the *E. heros* hybridization zone. However, soybean, the preferred host of *E. heros*, is also commonly grown in these areas during the first crop season. Thus, inferring the contribution of *E. heros* strains (pure and their hybrids) to dispersal to cotton-growing areas is crucial to understand this phenomenon and the evolutionary aspects of exploiting a new host. Therefore, our objectives were: 1) to test if there is a preference for a host (soybean or cotton) among the *E. heros* pure strains and their reciprocal hybrids ♀SS × ♂NS (hereafter HSN) and ♀NS × ♂SS (hereafter HNS), and 2) to evaluate the development and survivorship of nymphs of the two *E. heros* pure strains and their reciprocal hybrids on soybean and cotton. The evolutionary implications of the *E. heros* strains and the impacts on management of this pest are discussed.

## 2. Material and Methods

### 2.1 Collection and maintenance of *E. heros* stock colony strains

Individuals from the NS were collected in the municipality of Balsas, state of Maranhão (07°13′44.50′′ S, 45°58′35.32′′ W) and individuals from the SS were collected in Santa Maria, state of Rio Grande do Sul (29°42’17.9”S, 53°35’47.9”W). Approximately 200 adults from each locality were collected and taken to the Laboratory of Arthropod Molecular Ecology at the University of São Paulo, USP/ESALQ (Piracicaba, São Paulo, Brazil). The insects were reared and maintained as described by Hickmann et al. (2021). In brief, the insects were kept in cages under controlled laboratory conditions (25ºC, 65 % RH, and 14:10 (L:D) light regime) and natural diet and water were offered *ad libitum*.

Both *E. heros* strains were collected on soybean crops without nearby cotton crops, since cotton is grown mainly in central Brazil (CONAB, 2022). Both strains were maintained for approximately five generations without contact with soybean or cotton plants or introduction of field-collected insects before being used in the trials.

### 2.2 Cultivated plants

To obtain the reproductive structures and branches in the reproductive phase, seeds of soybean cv. “Brasmax Desafio RR” (approximately 30 plants per square meter) and cotton cv. “DP 1742 RF” (approximately 10 plants per square meter) were sown in the experimental area of the Department of Entomology and Acarology (Piracicaba, SP, Brazil) in October (cotton) and November (soybean) of 2020. The experimental area has red latosol, and we incorporated ≅ 8 g of fertilizer (4: 20: 20, N: P: K, respectively) per square meter before sowing.

### 2.3 Host-choice bioassays

We used individuals from populations of the two *E. heros* strains (SS and NS) established in the laboratory from the above localities to test for host preference. Reciprocal hybrids were obtained by crossing a ♀NS with a ♂SS = HNS and ♀SS with a ♂NS = HSN (15 couples for each cross). Host-choice tests were carried out with nymphs of the 4^th^ instar (two days after instar change) and adult females of reproductive age (≅ two weeks old) from both strains of *E. heros* and the reciprocal hybrids HNS and HSN. The insects were starved for 24 h before testing and water was offered *ad libitum*. The experiments consisted of 30 repetitions for each strain, reciprocal hybrid, and life stage.

The host-choice tests were performed in a dark room with controlled temperature and humidity (25 ± 1°C and 65 ± 10% RH). The choice tests were carried out in transparent acrylic cages measuring 40 × 20 × 20 cm (length, width, and height, respectively). The soybean host used in the experiments was at reproductive stages R_3_– R_4_ (Fehr et al., 1971); in each choice test, six pods in the grain-filling phase plus a leaf of the upper trefoil were inserted into one end of the cage. For the cotton host, plants in principal growth stage 7, (71–79) (Munger et al., 1998) were used; a developed cotton bud and a healthy cotton leaf were placed at the opposite end of the cage. In each repetition, the cage was cleaned with 70% ethanol, and the hosts were replaced and inserted in alternate ends of the cage to prevent bias. The fasting insect, nymph or adult female, was released into the center of the cage and could choose between the soybean and cotton hosts. Visual evaluations were made after 10, 20 and 30 min for nymphs and 20, 40 and 60 min for females, noting which host the insects chose. If after the maximum stipulated time a choice had not occurred, the insect was not counted.

### 2.4 Nymph development and survivorship of *E. heros* SS, NS, HSN, and HNS on soybean or cotton

To identify developmental differences between strains and reciprocal hybrids fed exclusively on soybean or cotton, 70 2nd-instar nymphs (the first instar does not feed and is gregarious) of SS, NS, HSN, and HNS were transferred to cylindrical PVC cages (29.5 cm high and 24.5 cm in diameter). Cages were attached to a Styrofoam plate and covered with voile fabric containing soybean or cotton branches arranged in 100-mL flasks with distilled water. The experiments consisted of three repetitions for each strain and reciprocal hybrid and host. The branches were replaced every two days. The development time (days), weight (mg), and survivorship of each stage were evaluated daily for the nymphs on both hosts.

### 2.5 Statistical analysis

Data from the host-choice bioassays were processed with the Fisher test and a chi-square test in R (R Core Team 2022). The biological data were tested for normality with the Shapiro-Wilk test in R (R Core Team 2022). Data that did not fit a normal distribution were normalized (log). Data were then submitted to a variance analysis and the means submitted to a Tukey test (p = 0.05). The survival over time of the immature phases was submitted to a Log-rank test in the R package “nph” (Ristl 2020).

## 3. Results

### 3.1 Host choice

The host-choice results showed significant differences in preference between soybean and cotton (Fisher’s Exact Test, p = 0.03852). SS nymphs and females showed a preference for the soybean host [Females: χ^*2*^ = 6.5333, df = 1, p = 0.01059; Nymphs: χ^*2*^ = 8.5333, df = 1, p = 0.003487] (Figure 1). On the other hand, NS nymphs and females [Females: χ^*2*^ = 0.53333, df = 1, p = 0.4652; Nymphs: χ^*2*^ = 0, df = 1, p = 1], HSN [Females: χ^*2*^ = 1.2, df = 1, p = 0.2733; Nymphs: χ^*2*^ = 1.2, df = 1, p = 0.2733] and, HNS [Females: χ^*2*^ = 0.53333, df = 1, p = 0.4652; Nymphs: χ^*2*^ = 0.53333, df = 1, p = 0.4652] did not show a preference, choosing randomly between soybean and cotton (Figure 1).

**Figure 1.**
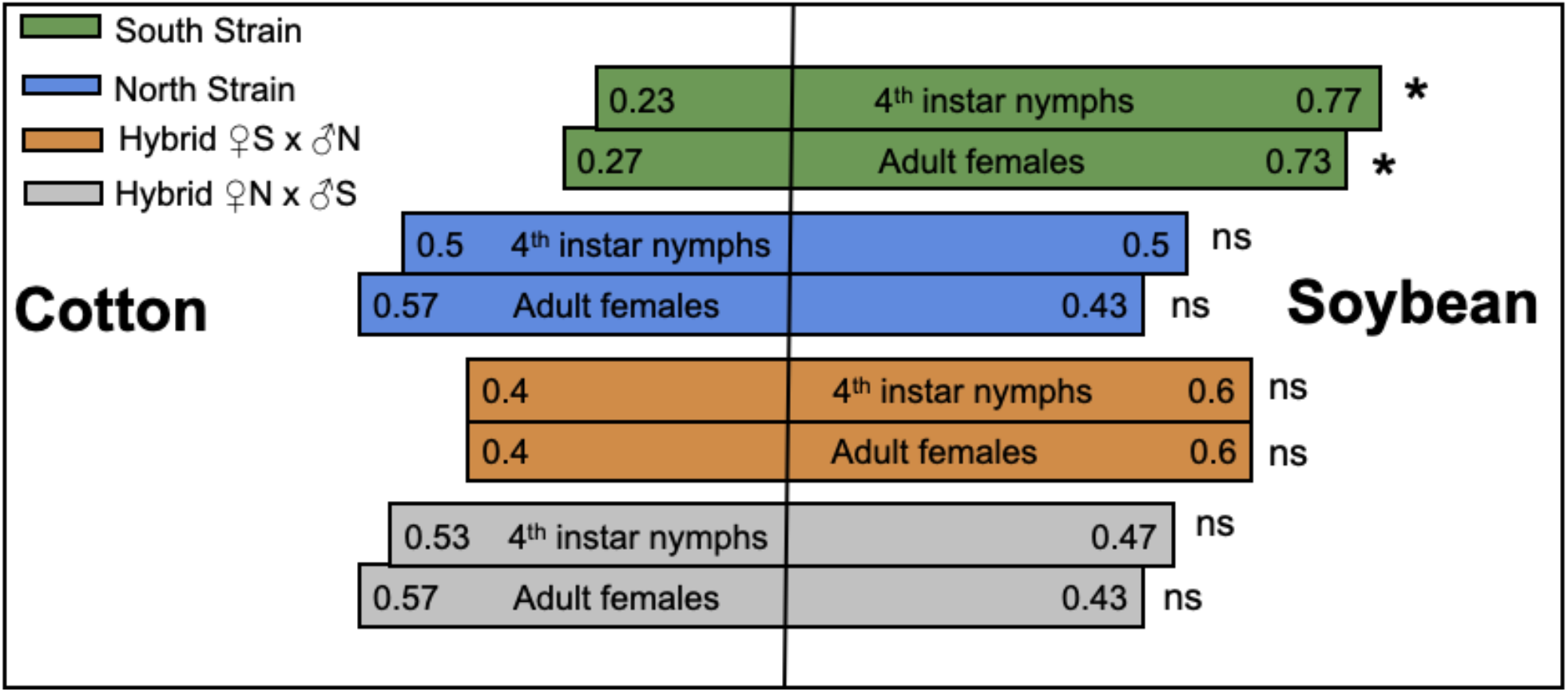
Choice proportion of *Euschistus heros* strains (south strain: SS and north strain: NS) and F1 hybrid offspring ♀N × ♂S (HNS) and ♀S × ♂N (HSN) between soybean and cotton plants. Asterisk indicates significant difference, and ns indicates lack of significance in a χ^2^ test (p<0.05).

### 3.2 Nymph development and survivorship of SS, NS, HSN, and HNS (hybrids) fed on soybean or cotton

#### Nymph development time

The development time of 2^nd^ instars differed between the hosts (soybean: 4.0 ± 0.5 to 5.7 ± 0.7 days and cotton: 7.3 ± 0.3 to 11.7 ± 0.3 days; F_1;7_ = 14.15, p<0.000) and between strains on cotton (NS: 11.7 days and HNS: 7.3 days; F_1;7_ = 14.15, p<0.000) (Table 1). The SS, NS, and HSN fed on cotton required more days to develop in the 2^nd^ instar than the HNS strain. We found no differences in the development time of 3^rd^-instar nymphs among the pure and hybrid strains when fed on soybean or cotton (F_1;7_ = 1.16, p< 0.377). Nymphs of the 4^th^ instar fed on soybean did not show differences among pure and hybrid strains (F_1;3_ = 7.75 p< 0.094); on the other hand, NS nymphs required a longer time to complete the 5^th^ instar on soybean (NS: 14.7 ± 0.5 F_1;3_ = 12.21, p = 0.002). Finally, we found no difference in the time needed to reach adulthood among the pure/hybrid strains on soybean, per Tukey test: SS: 34.0 ± 2.4; NS: 36.0 ± 4.0; HSN: 23.0 ± 1.2 and HNS: 23.0 ± 0.5 (F_1;3_ = 5.51, p = 0.0239) (Table 1).

**Tabela 1.**
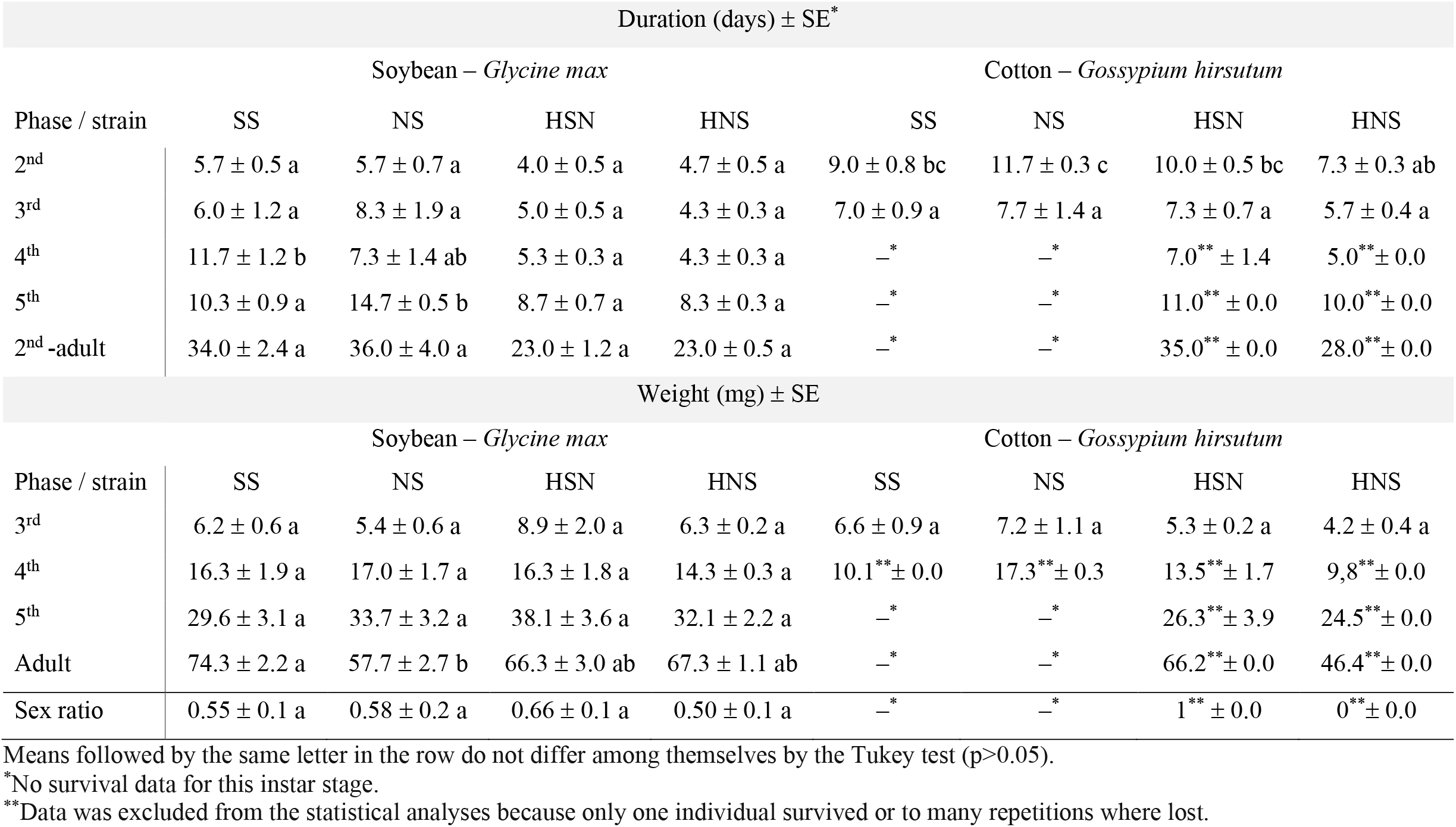
Biological parameters development time (mean days ± SE), weight (mean mg ± SE), sex ratio (mean ± SE) of *Euschistus heros* South Strain (SS), North Strain (NS), reciprocal hybrid ♀SS X ♂NS (HSN) and ♀NS X ♂SS (HNS) fed only on soybean or cotton under laboratory conditions (25 ± 1°C; 60 ± 10% RH; photophase 14 h).

#### Weight

The weight of the 3^rd^ instars did not differ among pure/hybrid strains when fed on cotton or soybean hosts, ranging from 4.2 ± 0.4 to 8.9 ± 2.0 mg (F_1;7_ =1.66, p = 0.1894; Table 1). Additionally, we found no difference in the weight of 4^th^ and 5^th^ instars among SS, NS, HSN, and HNS when fed on soybean (F_1;3_ = 0.36, p = 0.7826 and F_1;3_ = 0.90, p = 0.4817, 4^th^ and 5^th^ instars, respectively; Table 1). However, the weight of adults differed between SS (74.3 ± 2.2) and NS (57.7 ± 2.7); the hybrids showed an intermediate weight (F_1;3_ = 5.61, p = 0.0228; Table 1). The sex ratio of adults did not differ significantly among pure, or hybrid strains fed on soybean, ranging from 0.50 ± 0.1 to 0.66 ± 0.1 (F_1;3_ = 0.39, p = 0.7626; Table 1).

#### Nymph survivorship of SS, NS, HSN, and HNS on soybean or cotton

The average survivorship of pure/hybrid strains when fed exclusively with soybean or cotton showed a significant difference among pure/hybrid strains and host (Log-rank p = 0.0001 (Figure 2); confidence intervals are shown in Supplementary Table 1. Pure and hybrid strains of *E. heros* completed their life cycle on soybean (proportional values, SS: 0.07 ± .01, NS: 0.14 ± 0.06, HSN: 0.13 ± .00 and HNS: 0.24 ± 0.11; F_3;11_ = 1.5460, p = 0.2762; Figures 2 and 3). On the other hand, only the hybrid strains developed on cotton, although with low survivorship, less than 1% (SS: 0.0; NS:0.0; HSN: 0.003; HNS: 0.003; Figures 2 and 3).

**Figure 2.**
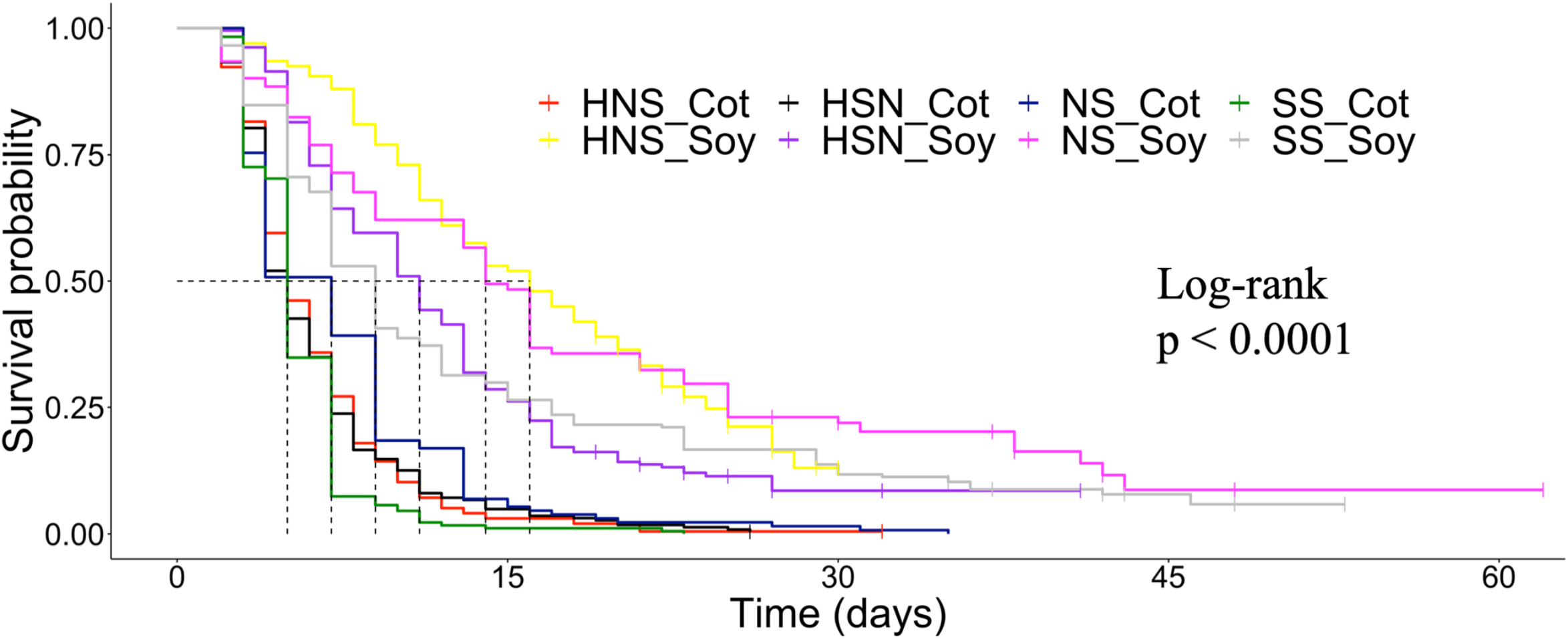
Median survival of South strain, North strain, reciprocal hybrid ♀N × ♂S (HNS), and reciprocal hybrid ♀S × ♂N (HSN) of *Euschistus heros* when fed on soybean or cotton differed significantly between strain/hybrid and host (Log-rank p = 0.0001). Red line: HNS fed only on cotton; yellow: HNS fed on soybean; black: HSN fed on cotton; purple: HSN fed on soybean; blue: NS fed on cotton; magenta: NS fed on soybean; green: SS fed on cotton; and gray: SS fed on soybean.

**Figure 3.**
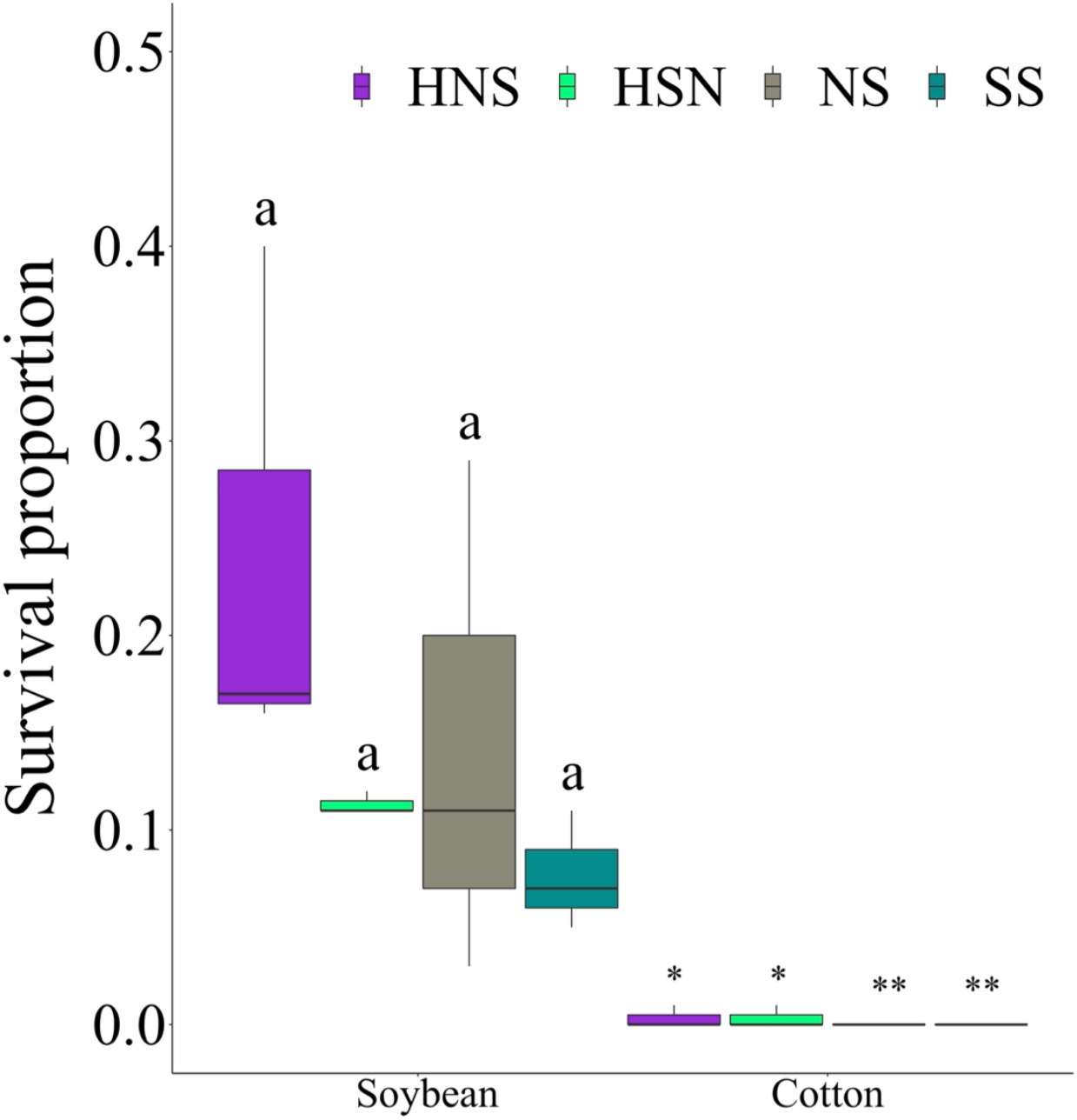
Survival proportion of *Euschistus heros* nymphs (2nd instar–adult) of South strain (SS), North strain (NS), and F1 hybrids (♀SS × ♂NS (HSN) and ♀NS × ♂SS (HNS)) fed on cotton or soybean plants. Boxplots with the same letter do not differ from each other by the Tukey test (p > 0.05). * Only one individual survived. ** all individuals died. (* and ** data were excluded from the statistical analyses because only one or no individual survived).

## 4. Discussion

The strains of *E. heros* showed preferences between soybean and cotton plants, depending on the strain. The SS strain chose soybean over cotton, suggesting a higher attraction to soybean plants, whereas the NS chose soybean and cotton evenly, showing a more polyphagous choice behavior. The reciprocal hybrid strains (HSN and HNS) behaved similarly to the NS, choosing randomly between cotton and soybean. Nymph survivorship of both the pure strains and their reciprocal hybrids was similar when fed on soybean. On the other hand, only reciprocal hybrids reached adulthood when fed on cotton, although with low viability. The suitability of the soybean host and the unsuitability of cotton for nymph development found for the *E. heros* strains has been related to the nutritional quality and allomone substances of the host plants (Azambuja et al., 2013; Panizzi, 2000).

The tightened relationship of the SS to the soybean host and the non-choice behavior of *E. heros* NS can be attributed to the historical and contemporary distribution of soybean and cotton crops in Brazil. After soybeans were introduced into Brazil in 1901, they were grown only in the southern region until 1980, when soybean cultivation began to expand to the central and north/northeastern regions (Cattelan and Dall’Agnol, 2018). Cotton-growing areas have a contrasting history of geographic expansion. Cotton has been cultivated for centuries in the Brazilian Northeast, and many perennial cotton cultivars are found in the north/northeastern regions. In the last century, cotton cultivation expanded to the southeast and central parts of the country (Barros et al., 2022). The historical expansion of soybean and cotton crops matches the occurrence areas of the allopatric strains of *E. heros* in Brazil and may account for the preference behavior of *E. heros* NS and SS.

In central Brazil, cotton fields are interspersed with soybean fields, which favors dispersal of insects from soybean to cotton (Soria et al 2010). *Euschistus heros* populations colonize soybean, and after soybean senescence the insects disperse to cotton plants, especially cotton bolls (Soria et al., 2017, 2011, 2010, 2009). In this scenario, *E. heros* may expand its host range because cotton becomes a resource for *E. heros* adults after soybean senescence. *Euschistus heros* is a multivoltine species with rare long-dispersal behavior (Panizzi, 1997; Soares et al., 2018). Thus, *E. heros* depletes local host-plant resources, or the host plant enters senescence, meaning that the insects must shift between host plants while seeking new resources in a limited spatial area (Carrasco et al., 2015; Tillman et al., 2009; Todd and Herzog, 1980; Venugopal et al., 2014).

The landscape changes experienced by *E. heros* populations in recent decades imply that the populations are under host-selection pressure, which may lead to host specialization or even ecological speciation (An et al., 2016; Funk, 2010; Zucchi et al., 2019). Host shift/expansion usually starts with recognizing the host. The non-choice preference between cotton and soybean by NS and hybrids suggests that *E. heros* may evolve to expand to cotton (Knolhoff and Heckel, 2014). Similarly, the preference of *E. heros* SS for soybean shows that this strain has adapted to this exotic host.

Both *E. heros* strains and their reciprocal hybrids completed their life cycle on soybean with similar biological parameters, in agreement with previous reports that soybean is a highly suitable host for *E. heros* (Azambuja et al., 2013; Possebom et al., 2020). However, the *E. heros* strains were inviable or showed low viability on cotton plants. When fed on cotton, the poor nymph survivorship indicates that the insect’s biology was significantly affected by the host, denoting a lower suitability of cotton. The faster adaptation of *E. heros* strains to the exotic soybean plants than to cotton is explained by the prevalence of species of Fabaceae as native hosts of *E. heros* (Link and Grazia, 1987; Panizzi and Lucini, 2017; Smaniotto and Panizzi, 2015). In contrast, no native Malvaceae species has been reported to support the complete development of *E. heros* (Azambuja et al., 2013; Panizzi and Lucini, 2017).

Although the reciprocal hybrids performed poorly on the cotton host, some nymphs did reach the adult phase. This observation is highly concerning, especially because a very large number of insects move annually from soybean to cotton fields. The recurrent attacks and colonization of cotton fields may be selecting insects with a higher capacity to colonize cotton plants (Joshi and Thompson, 1995). The suitability of an herbivore host is mediated by an insect’s capacity to metabolize the primary nutrients and secondary metabolites (allelochemicals) contained in the host (Schoonhoven et al., 1998a; War et al., 2012), suggesting that *E. heros* has mostly overcome the defenses of soybean and is still in the early steps of adapting to cotton (a gossypol-rich plant).

Besides the movement of *E. heros* adults into cotton fields when seeking for food, the ecology of this species must be considered. *Euschistus heros* adults show a facultative diapause triggered by short-day conditions, 12 h light or less per day (Mourão and Panizzi, 2002, 2000a, 2000b). The majority of Brazilian soybean and cotton fields are in latitudes above 8° South (CONAB 2022, IBGE 2022), meaning at least three months of short-day conditions per year. Therefore, after the soybean harvest, the adults may disperse to cotton fields, seeking shelter and food to survive the coming mild winter and not necessarily to develop on the host. Therefore, it is imperative to determine whether the *E. heros* insects that arrive in cotton fields are in diapause, and the contribution of dispersing insects from cotton to soybean fields in the next crop season. Furthermore, insecticides are applied to cotton more frequently than to soybean, with reports of 20–25 applications a year, with two applications for *E. heros* control in some areas (Lamas and Chitarra, 2014; Miranda, 2010; Pitta et al., 2018). The movement of *E. heros* individuals between cotton and soybean crops could accelerate the selection of *E. heros* populations by insecticides, raising serious concerns for management of this species (Tuelher et al., 2018).

Intra- and interpopulation variations in host use have been attributed to different plants in different regions or to genetic differences among populations (Carrière and Roitberg, 1995; Jaenike, 1990; Schoonhoven et al., 1998b). The agricultural expansion in Brazil has changed the landscape dramatically and has directly affected native insect communities, introducing novel selection pressures (Corrêa et al., 2019). For instance, rapid selection of crop pests for host use has been reported in other insect groups (Bernal and Medina, 2018; Gray et al., 2009). Here, we demonstrated differences among *E. heros* strains in host-choice responses and survivorship on soybean and cotton. *Euschistus heros* SS showed a higher preference for soybean, whereas NS and laboratory reciprocal hybrids showed a polyphagous choice behavior, suggesting that the selection process for host may have been mediated by historical and, mainly, the contemporary relationship of *E. heros* strains with these hosts.

In conclusion, we found a consistent host choice/survivorship pattern among *E. heros* strains in our bioassays. These different *E. heros* phenotypes characterized in the laboratory need to be tested in field conditions, to better understand the role of these strains (pure and hybrids) in the host-range adaptation. It is imperative to continue genotypic and phenotypic monitoring of *E. heros* from different agricultural landscapes, since the changes in host ranges would affect other ecological traits such as insecticide susceptibility, overwintering ecology, and interactions with natural enemies.

## References

An, X.-K., Sun, L., Liu, H.-W., Liu, D.-F., Ding, Y.-X., Li, L.-M., Zhang, Y.-J., Guo, Y.-Y., 2016. Identification and expression analysis of an olfactory receptor gene family in green plant bug Apolygus lucorum (Meyer-Dür). Sci. Rep. 6, 37870. https://doi.org/10.1038/srep37870

Azambuja, R., Degrande, P.E., Pereira, F.F., 2013. Comparative biology of Euschistus heros (F.) (Hemiptera: Pentatomidae) feeding on cotton and soybean reproductive structures. Neotrop. Entomol. 42, 359–365. https://doi.org/10.1007/s13744-013-0132-6

Barman, A.K., Parajulee, M.N., Sansone, C.G., Medina, R.F., 2012. Host preference of cotton fleahopper, Pseudatomoscelis seriatus (Reuter) is not labile to geographic origin and prior experience. Environ. Entomol. 41, 125–132. https://doi.org/10.1603/EN11221

Barros, M.A.L., Silva, C.R.C. Da, Lima, L.M. De, Farias, F.J.C., Ramos, G.A., Santos, R.C. Dos, 2022. A review on evolution of cotton in Brazil: GM, white, and colored cultivars. J. Nat. Fibers 19, 209–221. https://doi.org/10.1080/15440478.2020.1738306

Bernal, J.S., Medina, R.F., 2018. Agriculture sows pests: how crop domestication, host shifts, and agricultural intensification can create insect pests from herbivores. Curr. Opin. Insect Sci. 26, 76–81. https://doi.org/10.1016/j.cois.2018.01.008

Cahenzli, F., Erhardt, A., 2013. Transgenerational acclimatization in an herbivore–host plant relationship. Proc. R. Soc. B Biol. Sci. 280, 20122856. https://doi.org/10.1098/rspb.2012.2856

Carrasco, D., Larsson, M.C., Anderson, P., 2015. Insect host plant selection in complex environments. Curr. Opin. Insect Sci. 8, 1–7. https://doi.org/10.1016/j.cois.2015.01.014

Carrière, Y., Roitberg, D., 1995. Evolution of host-selection behaviour in insect herbivores: genetic variation and covariation in host acceptance within and between populations of Choristoneura rosaceana (Family: Tortricidae), the obliquebanded leadfoller. Heredity. 74, 357–368. https://doi.org/10.1038/hdy.1995.54

Cattelan, A.J., Dall’Agnol, A., 2018. The rapid soybean growth in Brazil. OCL 25, 1–12. https://doi.org/10.1051/ocl/2017058

CONAB, 2022. Acompanhamento da Safra Brasileira de Grãos, Boletim de Acompanhamento da Safra Brasileira de Grãos. Brasilia. https://www.conab.gov.br/info-agro/safras/graos/boletim-da-safra-de-graos. (accessed 24 May 2022).

Corrêa, A.S., Cordeiro, E.M., Omoto, C., 2019. Agricultural insect hybridization and implications for pest management. Pest Manag. Sci. 75, 2857–2864. https://doi.org/10.1002/ps.5495

Dávila-Flores, A.M., DeWitt, T.J., Bernal, J.S., 2013. Facilitated by nature and agriculture: performance of a specialist herbivore improves with host-plant life history evolution, domestication, and breeding. Oecologia 173, 1425–1437. https://doi.org/10.1007/s00442-013-2728-2

Fehr, W.R., Caviness, C.E., Burmood, D.T., Pennington, J.S., 1971. Stage of development descriptions for soybeans, Glycine Max (L.) Merrill 1. Crop Sci. 11, 929–931. https://doi.org/10.2135/cropsci1971.0011183X001100060051x

Funk, D.J., 2010. Does strong selection promote host specialisation and ecological speciation in insect herbivores? Evidence from Neochlamisus leaf beetles. Ecol. Entomol. 35, 41–53. https://doi.org/10.1111/j.1365-2311.2009.01140.x

Gasperi, G., Guglielmino, C.R., Malacrida, A.R., Milani, R., 1991. Genetic variability and gene flow in geographical populations of Ceratitis capitata (Wied.) (medfly). Heredity. 67, 347–356. https://doi.org/10.1038/hdy.1991.98

Gray, M.E., Sappington, T.W., Miller, N.J., Moeser, J., Bohn, M.O., 2009. Adaptation and invasiveness of western corn rootworm: intensifying research on a worsening pest. Annu. Rev. Entomol. 54, 303–321. https://doi.org/10.1146/annurev.ento.54.110807.090434

Hickmann, F., Cordeiro, E.G., Soares, P.L., Aurélio, M.S.L., Schwertner, C.F., Corrêa, A.S., 2021. Reproductive patterns drive the gene flow and spatial dispersal of Euschistus heros (Hemiptera: Pentatomidae). J. Econ. Entomol. 114, 2346–2354. https://doi.org/10.1093/jee/toab190

IBGE – Brazilian Institute of Geography and Statistics. https://www.ibge.gov.br/estatisticas/economicas/agricultura-e-pecuaria/9201-levantamento-sistematico-da-producao-agricola.html?=&t=destaques. (accessed 24 May 2022).

Jaenike, J., 1990. Host specialization in phytophagous insects. Annu. Rev. Ecol. Syst. 21, 243–273. https://doi.org/10.1146/annurev.es.21.110190.001331

Joshi, A., Thompson, J.N., 1995. Trade-offs and the evolution of host specialization. Evol. Ecol. 9, 82–92. https://doi.org/10.1007/BF01237699

Knolhoff, L.M., Heckel, D.G., 2014. Behavioral assays for studies of host plant choice and adaptation in herbivorous insects. Annu. Rev. Entomol. 59, 263–278. https://doi.org/10.1146/annurev-ento-011613-161945

Lamas, F.M., Chitarra, L.G., 2014. Diagnóstico dos Sistemas de Produção de Algodão em Mato Grosso. Embrapa Agropecuária Oeste Dourados, MS, 2014, 32p.

Link, D., Grazia, J., 1987. Pentatomídeos da região central do Rio Grande do Sul (Heteroptera). An. da Soc. Entomológica Bras. do Bras. 16, 115–129.

Medina, R.F., Reyna, S.M., Bernal, J.S., 2012. Population genetic structure of a specialist leafhopper on Zea: likely anthropogenic and ecological determinants of gene flow. Entomol. Exp. Appl. 142, 223–235. https://doi.org/10.1111/j.1570-7458.2012.01220.x

Miranda, J.E., 2010. Manejo Integrado de Pragas do Algodoeiro no Cerrado Brasileiro. Embrapa - Circ. Técnica 36. Campina Grande, PB, 2010, 37p.

Mourão, A.P.M., Panizzi, A.R., 2002. Photophase influence on the reproductive diapause, seasonal morphs, and feeding activity of Euschistus heros (Fabr., 1798) (Hemiptera: Pentatomidae). Brazilian J. Biol. 62, 231–238. https://doi.org/10.1590/S1519-69842002000200006

Mourão, A.P.M., Panizzi, A.R., 2000a. Estágios ninfais fotossensíveis à indução da diapausa em Euschistus heros (Fabr.) (Hemiptera: Pentatomidae). An. da Soc. Entomológica do Bras. 29, 219–225. https://doi.org/10.1590/S0301-80592000000200003

Mourão, A.P.M., Panizzi, A.R., 2000b. Diapausa e diferentes formas sazonais em Euschistus heros (Fabr.) (Hemiptera : Pentatomidae) no Norte do Paraná. An. da Soc. Entomológica do Bras. 29, 205–218. https://doi.org/10.1590/S0301-80592000000200002

Munger, P., Bleiholder, H., Hack, H., Hess, M., Stauβ, R., Boom, T., Weber, E., 1998. Phenological growth stages of the cotton plant (Gossypium hirsutum L.): codification and description according to the BBCH scale. J. Agron. Crop Sci. 180, 143–149. https://doi.org/10.1111/j.1439-037X.1998.tb00384.x

Panizzi, A. E. McPherson, J., James, D., Javahery, M., M. McPherson, R., 2000. Economic Importance of stink bugs (Pentatomidae) Heteroptera of Economic Importance, CRC Press, Boca Raton, FL., USA, in: Schaefer, C.W., Panizzi, A.. (Eds.), Heteroptera of Economic Importance. CRC Press, Boca Raton, FL, pp. 421–474.

Panizzi, A.R., 2000. Suboptimal nutrition and feeding behavior of hemipterans on less preferred plant food sources. An. da Soc. Entomológica do Bras. 29, 1–12. https://doi.org/10.1590/S0301-80592000000100001

Panizzi, A.R., 1997. Wild hosts of pentatomids: ecological significance and role in their pest status on crops. Annu. Rev. Entomol. 42, 99–122. https://doi.org/10.1146/annurev.ento.42.1.99

Panizzi, A.R., Lucini, T., 2017. Host plant-stinkbug (Pentatomidae) relationships, in: Čokl, A., Borges, M. (Eds.), Stinkbugs: Biorational Control Based on Communication Processes. CRC Press Taylor & Francis Group, Boca Raton, FL, pp. 31–58.

Panizzi, A.R., Slansky, F. Jr., 1985. Review of phytophagous pentatomids (Hemiptera: Pentatomidae) associated with soybean in the Americas. Florida Entomol. 68, 184–214.

Pitta, R.M., Rodrigues, S.M.M., Vivan, L.M., Bianchin, K.A., 2018. Susceptibility of Euschistus heros (Fabr 1794.) (Heteroptera: Pentatomidae) to insecticides in Mato Grosso. Sci. Electron. Arch. 11, 1–5.

Possebom, T., Lucini, T., Panizzi, A.R., 2020. Stink bugs nymph and adult biology and adult preference on cultivated crop plants in the southern Brazilian Neotropics. Environ. Entomol. 49, 132–140. https://doi.org/10.1093/ee/nvz142

Ristl, R. 2020. nph: planning and analyzing survival studies under non-proportional hazards. R package version 2.0. https://CRAN.Rproject.org/package=nph

Schoonhoven, L.M., Jermy, T., van Loon, J.J.A., 1998a. Plants as insect food: not the ideal, in: Schoonhoven, L.M., Jermy, T., van Loon, J.J.A. (Eds.), Insect-Plant Biology: From Physiology to Evolution. Chapman and Hall, London, pp. 83–113.

Schoonhoven, L.M., Jermy, T., van Loon, J.J.A., 1998b. Host-plant selection: why insects do not behave normally, in: Schoonhoven, L.M., Jermy, T., van Loon, J.J.A. (Eds.), Insect-Plant Biology: From Physiology to Evolution. Chapman and Hall, London, pp. 195–214.

Silva, C.S., Cordeiro, E.M.G., Paiva, J.B., Dourado, P.M., Carvalho, R.A., Head, G., Martinelli, S., Correa, A.S., 2020. Population expansion and genomic adaptation to agricultural environments of the soybean looper, Chrysodeixis includens. Evol. Appl. eva.12966. https://doi.org/10.1111/eva.12966

Simon, J.-C., D’Alencon, E., Guy, E., Jacquin-Joly, E., Jaquiery, J., Nouhaud, P., Peccoud, J., Sugio, A., Streiff, R., 2015. Genomics of adaptation to host-plants in herbivorous insects. Brief. Funct. Genomics 14, 413–423. https://doi.org/10.1093/bfgp/elv015

Smaniotto, L.F., Panizzi, A.R., 2015. Interactions of selected species of stink sugs (Hemiptera: Heteroptera: Pentatomidae) from leguminous crops with plants in the Neotropics. Florida Entomol. 98, 7–17. https://doi.org/10.1653/024.098.0103

Soares, P.L., Cordeiro, E.M.G., Santos, F.N.S., Omoto, C., Correa, A.S., 2018. The reunion of two lineages of the Neotropical brown stink bug on soybean lands in the heart of Brazil. Sci. Rep. 8, 2496. https://doi.org/10.1038/s41598-018-20187-6

Soria, M.F., Degrande, P.E., Panizzi, A.R., 2011. Symptoms, injuries, yield reduction and quality loss of cotton attacked by the neotropical brown stink bug Euschistus heros (F.) (Hemiptera: Pentatomidae), in: Beltwide Cotton Conferences. Atlanta-GA, pp. 4–7.

Soria, M.F., Degrande, P.E., Panizzi, A.R., 2010. Algodoeiro invadido. Cultivar 18–20.

Soria, M.F., Degrande, P.E., Panizzi, A.R., Toews, M.D., 2017. Economic injury level of the Neotropical brown stink bug Euschistus heros (F.) on cotton plants. Neotrop. Entomol. 46, 324–335. https://doi.org/10.1007/s13744-016-0454-2

Soria, M.F., Thomazoni, D., Rosa, R., Paulo, M., Degrande, E., 2009. Stink bugs incidence on bt cotton in Brazil, in: Beltwide Cotton Conferences. pp. 813–819.

Sosa-Gómez, D.R., Corrêa-Ferreira, B.S., Kraemer, B., Pasini, A., Husch, P.E., Delfino Vieira, C.E., Reis Martinez, C.B., Negrão Lopes, I.O., 2020. Prevalence, damage, management and insecticide resistance of stink bug populations (Hemiptera: Pentatomidae) in commodity crops. Agric. For. Entomol. 22, 99–118. https://doi.org/10.1111/afe.12366

Tillman, P.G., Northfield, T.D., Mizell, R.F., Riddle, T.C., 2009. Spatiotemporal patterns and dispersal of stink bugs (Heteroptera: Pentatomidae) in peanut-cotton farmscapes. Environ. Entomol. 38, 1038–1052. https://doi.org/10.1603/022.038.0411

Todd, J.W., Herzog, D.C., 1980. Sampling phytophagous Pentatomidae on soybean, in: Kogan, M., Herzog, D.C. (Eds.), Sampling Methods in Soybean Entomology. Springer-Verlang, New York, pp. 438–478.

Tuelher, E.S., da Silva, É.H., Rodrigues, H.S., Hirose, E., Guedes, R.N.C., Oliveira, E.E., 2018. Area-wide spatial survey of the likelihood of insecticide control failure in the neotropical brown stink bug Euschistus heros. J. Pest Sci. (2004). 91, 849–859. https://doi.org/10.1007/s10340-017-0949-6

Venugopal, P.D., Coffey, P.L., Dively, G.P., Lamp, W.O., 2014. Adjacent habitat influence on stink bug (Hemiptera: Pentatomidae) densities and the associated damage at field corn and soybean edges. PLoS One 9, e109917. https://doi.org/10.1371/journal.pone.0109917

War, A.R., Paulraj, M.G., Ahmad, T., Buhroo, A.A., Hussain, B., Ignacimuthu, S., Sharma, H.C., 2012. Mechanisms of plant defense against insect herbivores. Plant Signal. Behav. 7, 1306–1320. https://doi.org/10.4161/psb.21663

Wetzel, W.C., Kharouba, H.M., Robinson, M., Holyoak, M., Karban, R., 2016. Variability in plant nutrients reduces insect herbivore performance. Nature 539, 425–427. https://doi.org/10.1038/nature20140

Wilding, C.S., Butlin, R.K., Grahame, J., 2001. Differential gene exchange between parapatric morphs of Littorina saxatilis detected using AFLP markers. J. Evol. Biol. 14, 611–619. https://doi.org/10.1046/j.1420-9101.2001.00304.x

Zucchi, M.I., Cordeiro, E.M.G., Wu, X., Lamana, L.M., Brown, P.J., Manjunatha, S., Viana, J.P.G., Omoto, C., Pinheiro, J.B., Clough, S.J., 2019. Population genomics of the Neotropical brown stink bug, Euschistus heros: the most important emerging insect pest to soybean in Brazil. Front. Genet. 10, 1–12. https://doi.org/10.3389/fgene.2019.01035

