## Supplementary Table 1 for "Host preference and survivorship of *Euschistus heros* (Hemiptera: Pentatomidae) strains on cotton and soybean"

**Supplementary Table 1.** Survival means, standard error means, median and confidence intervals of nymph survival of *Euschistus heros* south strain (SS), north strain (NS) and reciprocal hybrids (♀N × ♂S (HNS) and ♀S × ♂N (HSN)) fed on soybean or cotton in laboratory conditions.

| | Records | n.max | n.start | events | r.mean | se(r.mean) | median | 0.95LCL | 0.95UCL |
|---|---|---|---|---|---|---|---|---|---|
| **SS Cotton** | 210 | 210 | 210 | 210 | 5.266667 | 1.0830551 | 5 | 5 | 5 |
| **SS Soybean** | 210 | 210 | 210 | 193 | 14.409524 | 0.1817693 | 9 | 7 | 9 |
| **NS Cotton** | 210 | 210 | 210 | 210 | 6.590476 | 0.3149625 | 4 | 4 | 7 |
| **NS Soybean** | 210 | 210 | 210 | 182 | 18.043999 | 1.2649172 | 13 | 9 | 15 |
| **HSN Cotton** | 210 | 210 | 210 | 209 | 6.623810 | 0.3870137 | 5 | 4 | 6 |
| **HSN Soybean** | 210 | 210 | 210 | 186 | 15.071977 | 1.3297792 | 11 | 10 | 12 |
| **HNS Cotton** | 210 | 210 | 210 | 209 | 6.214286 | 0.3567731 | 5 | 5 | 6 |
| **HNS Soybean** | 210 | 210 | 210 | 159 | 20.230694 | 1.6475465 | 15 | 13 | 18 |
